# ViPal: A Framework for Virulence Prediction of Influenza Viruses with Prior Viral Knowledge Using Genomic Sequences

**DOI:** 10.1101/2022.03.24.485635

**Authors:** Rui Yin, Zihan Luo, Pei Zhuang, Chee Keong Kwoh, Zhuoyi Lin

**Affiliations:** School of Computer Science and Engineering, Nanyang Technological University, 50 Nanyang Avenue, Singapore; Department of Biomedical Informatics, Harvard Medical School, Boston, USA; School of Computer Science and Technology, Huazhong University of Science and Technology, China; Department of Pharmaceutics, University of Florida, Gainesville, Florida, USA

## Abstract

Influenza viruses pose significant threats to public health and cause enormous economic loss every year. Previous work has revealed the viral factors that influence the virulence of influenza viruses. However, taking prior viral knowledge represented by heterogeneous categorical and discrete information into account is scarce in the existing work. How to make full use of the preceding domain knowledge into virulence study is challenging but beneficial. This paper proposes a general framework named ViPal for virulence prediction that incorporates discrete prior viral mutation and reassortment information based on all eight influenza segments. The posterior regularization technique is leveraged to transform prior viral knowledge to constraint features and integrated into the machine learning models. Experimental results on influenza genomic datasets validate that our proposed framework can improve virulence prediction performance over baselines. The comparison between ViPal and other existing methods shows the computational efficiency of our framework with superior performance. Moreover, the interpretable analysis through SHAP identifies the scores of constraint features contributing to the prediction. We hope this framework could provide assistance for the accurate detection of influenza virulence and facilitate flu surveillance.

## 1 Introduction

Influenza A virus is a member of the Orthomyxoviridae family that consists of eight single-stranded, negative-sense RNA segments, encoding up to 16 classic proteins. Among all the segments, hemagglutinin (HA) and neuraminidase (NA) are the most important that characterize influenza A viruses into distinct categories [24]. The HA segment is responsible for binding the virus to cells with sialic acid on the membranes [40]. It is initially synthesized as a single polypeptide precursor (HA0), which needs to be cleaved into subunits HA1 and HA2 by cellular proteases to become biologically active [2]. NA segment functions as a tetramer that cleaves sialic acid from cells and virion glycoproteins to prevent clumping of released viruses [3]. Up to now, 18 hemagglutinin sub-types (H1 to H18) and 11 neuraminidase subtypes (N1 to N11) have been identified [43]. Apart from the HA and NA segments, the influenza A virus particle comprises nucleoprotein (NP), non-structural proteins (NS1 and NS2), matrix proteins (M1 and M2), and three RNA polymerase subunits, polymerase acidic protein (PA), polymerase basic protein 1 (PB1) and polymerase basic protein 2 (PB2) [53]. The PA segment contains a second open reading frame that encodes the PA-X protein [43], while the PB1 segment can encode two more proteins PB1-F2 and PB1 N40. The RNA polymerase complex plays crucial roles in both transcription and replication of the viral genomes [7]. Similarly, the M1 protein mediates virion assembly and M2 protein forms the channels for the viral entry. Moreover, NS1 protein functions as an antagonist that inhibits interferon related activities [14] and NS2 protein has been implicated in mediating the nuclear export of ribonucleoprotein (RNP) complexes and recruiting ATPase for efficient viral exit [36].

Mutation and reassortment are the key mechanisms driving the evolution of influenza viruses and significantly increases the diversity of circulating strains. They have enabled changes in viral proteins and allowed for the exchange of the segments from different influenza strains to co-infect the host cells [28]. The emergence of novel strains may affect the virulence of viruses. The mutations in viral genes associated with virulence have been explored in many ways. Evidence showed that the mutations in the region 130-loop, 190-helix and 220-loop on HA protein increased its virulence during the adaptation of influenza A virus in mice [20]. Several potential virulent sites were identified in HA segment through rule-based methods using past pandemics strains [66]. The substitutions of E627K and D701N in PB2 segment have also been considered as general markers that played a significant role in the increased virulence in mice [22]. In addition to HA and PB2, the genetic markers from other segments can also influence virulence and pathogenicity. The mutations at position 223 and 275 in the NA protein [6, 27], 97 in PA [51] and 92 in NS1 [44] were related to enhanced virulence in mammalian hosts. Furthermore, it was discovered that the mutations at multiple sites on different genes would have a synergistic effect on influenza viruses’ virulence. For instance, the synergistic effect of dual mutations N383D and S224P in PA led to the increased polymerase activity and has been used as the hallmark for natural adaption of H1N1 and H5N1 viruses to mammals [50]. Another example was the synergistic action of two mutations D222G and K163E in HA protein with the mutation F35L in PA of 2009 H1N1 pandemic strain that causes lethality in the infected mouse [22, 45].

Compared with the influence on the virulence caused by mutations, recent studies of viral evolutionary dynamics have displayed the importance of reassortment in generating highly pathogenic epidemic and pandemic strains [33, 52]. The evidence suggested that the 1918 Spanish flu pandemic caused by influenza A H1N1 virus is a mixed recombination between the North American H1N2 swine influenza virus, European H1N1 swine influenza virus, North American avian influenza virus and H3N2 influenza virus [55]. Meanwhile, three other influenza pandemics were also associated with emerging strains due to the reassortment process. The reassortant strains from the combination of avian and human viruses are responsible for the two pandemics in 1957 and 1968, respectively [29, 47, 59]. It was indicated that the 1957 Asian flu originated from the 1918 Spanish flu. The HA and NA segments were derived from other avian viruses, while five of the rest of the segments were retained. Similarly, the reassortment of human and avian strains with an H3 HA gene stemmed from the avian virus led to the 1968 H3N2 pandemic [39]. The most recent pandemics in 2009 was due to a reassortant strain generated through the reassortment of human, avian and swine strains [48]. These pandemics not only engendered numerous deaths but also deteriorated the social economy. Furthermore, except for the pandemic strains, it was reported with human cases infected by the new emerging H7N9 strains [5]. These H7N9 strains were uncovered as progeny through the reassortment between H7N9 and H9N2 subtypes [13], which has the potential to trigger another flu pandemic. Therefore, the reassortment mechanism has significantly impacted the occurrence of novel strains with high virulence, potentially leading to the outbreak of pandemics that impacts a huge amount of people with illness and deaths. Deep neural networks (DNNs) have been widely applied in the field of bioinformatics for learning patterns from massive data, including mutation prediction [63], antigenicity estimation [70] [64], human microbe-drug association [30], protein secondary structure prediction [58], vaccine selection [65] and protein-protein interaction prediction [69], etc. Despite the significant progress in the performance of computational models, DNN methods still have limitations. The high predictive results have heavily relied on large amounts of training data. The purely deep learning neural networks have created a black box that leads to uninterpreted and sometimes counter-intuitive results [54, 34]. The cognitive process of humans indicates that people learn from not only concrete samples but also different types of experience and knowledge [26], such as the relationship between risk factors and diseases, the mutations on the influence of viral pathogenicity. As we know, prior biomedical knowledge could provide critical information that facilitates disease detection and diagnosis. However, it is easy to ignore the importance of general information and domain knowledge to construct computational models and regulate the learning process.

In this work, we propose a novel predictive framework that can successfully incorporate discrete prior viral knowledge into the models to improve the performance. This framework enables neural networks to learn simultaneously from label genomic sequences combined with corresponding prior viral knowledge, specifically mutation and reassortment information, to predict the virulence of influenza A viruses. This is through an iterative rule knowledge distillation process [17] that converts the encoded discrete prior knowledge into the network parameters. At each iteration, posterior regularization [12] is used to convert the prior genomic information by converting the posterior distribution into the constraint feature sets. The workflow of the proposed framework is shown in Figure 1. We conduct a variety of experiments on the collected influenza dataset. The results indicate that the proposed framework is capable of integrating prior viral knowledge and achieves better or comparable performance over the baseline and existing prediction models. The encouraging predictive results indicate that this model can be used to estimate the lethality of viruses and facilitate flu surveillance of the novel emerging influenza strains. The main contribution of the proposed framework is highlighted as follows:

**Figure 1:**
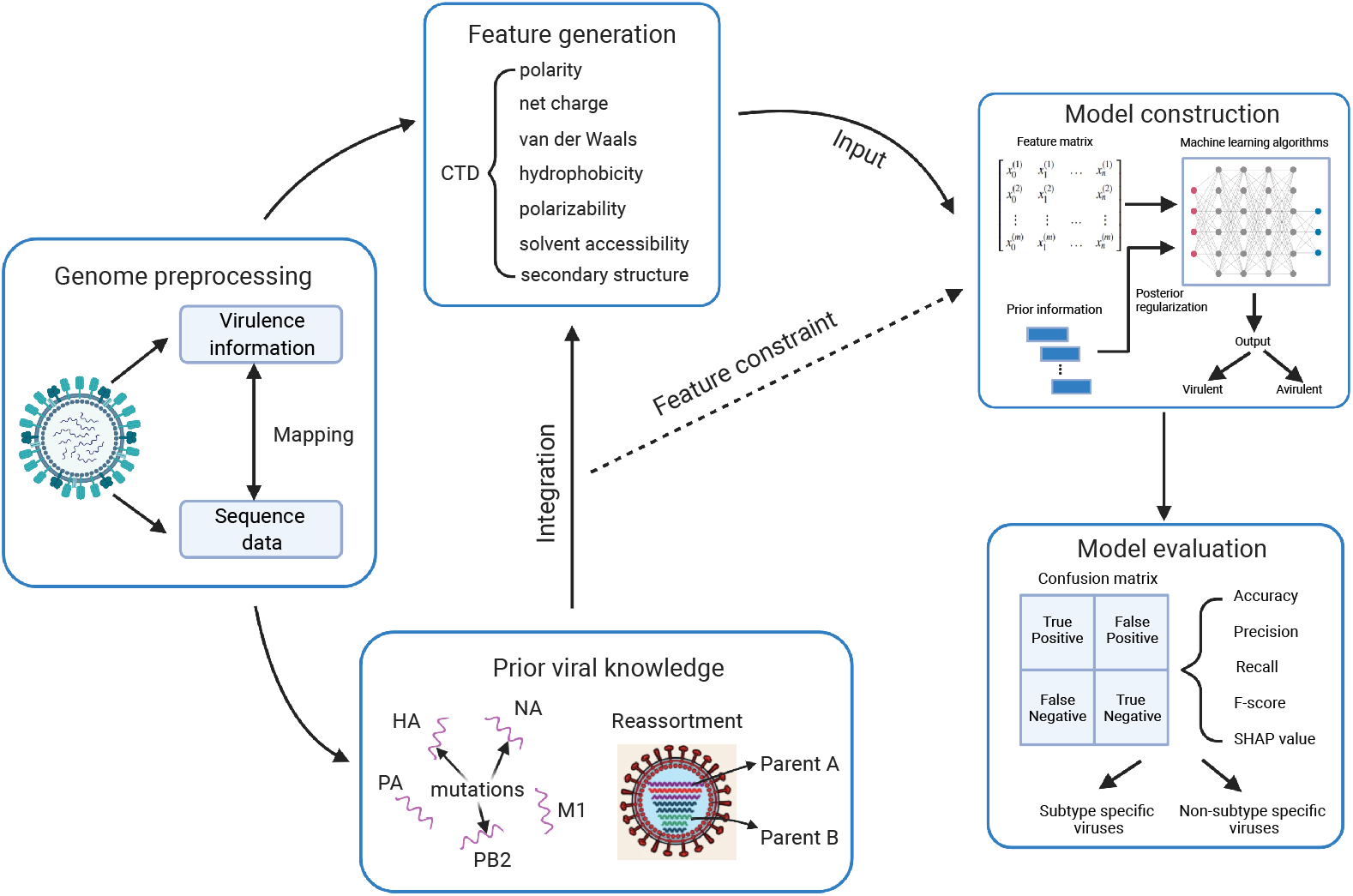
The workflow of proposed framework for virulence prediction integrating prior viral knowledge based on genomic data.

- This is the first time to integrate viral mutation and reassortment information as prior knowledge for virulence prediction.
- This framework ViPal utilizes posterior regularization and converts posterior distribution for the feature constraint.
- It is capable of interpreting the importance of different mutation sites and reassortment information of the predictive models.
- The results demonstrate superior effectiveness and computational efficiency for the task of influenza virulence prediction. Moreover, It is a general model and can be easily applied to other viruses.

## 2 Materials and methods

### 2.1 Problem formulation

Virulence usually refers to a pathogen’s ability to infect a resistant host in the context of gene systems [41], and yet there is no unified and rigorous definition of virulence. In this paper, we only explored the virulence of influenza A viruses. We followed the settings in [21] to measure the virulence of infections through mouse lethal dose (MLD) 50. The level of virulence is categorized into avirulent and virulent types, respectively. Each unique infection and its corresponding strain with known MLD50 value was collected. When the MLD50 is greater than 10E6.0, it is regarded as avirulent. Otherwise, it is virulent. If the samples can not be determined by the lower or upper bound of MLD50, RULE 1 to 6 from [21] are applied to provide the virulence class of infection. These procedures assist the selection of influenza strains with virulence information for data collection.

### 2.2 Data collection

The virulence information of influenza sequences was collected through previous publications and experiments. The genomic sequences of all viral segments were downloaded from the National Center for Biotechnology Information (NCBI) [42], while the MLD50 values were obtained from the literature information. If the genome of a virus was incomplete, basic local alignment search tool (BLAST) [1] was performed to search the most similar strains to supplement the genomic data. The collection ended up with 488 unique records of influenza A virus information with corresponding complete genome strains. The influenza datasets used in this study are available in Supplementary Materials S1.

### 2.3 Feature transformation

The sequences from the same virus were concatenated using BioEdit [15] with virulence label. To address the problem of different lengths for these concatenated genomes, we introduced three global descriptors [8], i.e., composition (C), transition (T) and distribution (D) that can encode length-free genomic sequences simultaneously. It was demonstrated to be efficient for host tropism prediction with high performance [68]. Next, seven physicochemical properties [9] (net charge, hydrophobicity, polarizability, normalized van der Waals, volume polarity, solvent accessibility and secondary structure) of amino acids were used to convert sequences into numerical vectors using AAindex [23]. Each amino acid was divided into three different groups in terms of the physicochemical properties [57] (Supplementary Materials S2). In combination with CTD descriptors formulated below, we finally obtained a 147-dimensional vector for each genome sample.

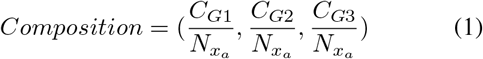

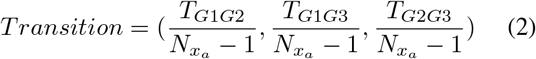

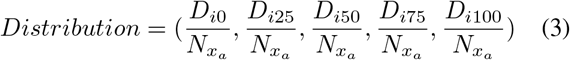

In Eq. (1), composition represents the percent frequency of amino acids of a particular property divided by the total number of amino acids. 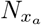 is the total number of amino acids from the input genome *x*_*a*_ and *C*_*Gi*_ is the frequency of amino acid property of group *i* in the genome. Transition in Eq. (2) characterizes the percentage frequency with which amino acid from a group is followed by another group denoted as *T*_*GiGj*_. It means the property in group *i* is followed by group *j* or the other way around such that *i, j* = 1,2,3 and *G*_*i*_ = *G*_*j*_. The third descriptor in Eq. (3) measures the distribution of each attribute in the sequence and *D*_*i*_ denotes the percentage in the positions of amino acid properties in group *i*. The distribution is based on the first, 25%, 50%, 75% and 100% of the amino acids for each attribute.

### 2.4 Constraint feature design

The measurement of influenza virulence remains a complicated and challenging issue. Viral mutation and reassortment are usually taken to investigate whether playing a role in the virulence change. The mutation information is at the sequence level, while the reassortment knowledge provides genomic information of the virus. Here we combine the prior viral knowledge into ViPal for the prediction of influenza virulence. In the subsequent section, we provide the design of the constraint features formulation.

#### Mutation information

It is natural to take mutation information into account for the exploration of influenza virulence. These mutations may occur at different sites on all influenza proteins. Table 1 summarizes some important mutations associated with virulence change or critical viral activities. We can observe that almost half of the mutations lie on HA protein that further indicates the importance of HA segment, e.g., mutation K163E increased the lethality in the infected mice and D185S is associated with pandemics. Besides, the mutations in other proteins also play crucial roles that make an impact on influenza virulence. Therefore, it is beneficial to incorporate prior mutation information as constraint features for constructing the prediction model.

**Table 1:** Literature summary of the molecular mutation markers of influenza viruses that are associated with virulence or critical viral activities.

| Protein | Position | Mutation | Determinant | References |
| --- | --- | --- | --- | --- |
| HA<sup>1</sup> | 138 | Ser (S) - Ala (A) | Sialic acid linkage | [61] |
|  | 143 | Ser (S) - Gly (G) | Reduced antigenicity in the hemagglutination inhibition test | [56] |
|  | 163 | Lys (K) - Glu (E) | Lethality in the infected mice | [45] |
|  | 185 | Asp (D) - Ser (S) | Associated with pandemics | [66] |
|  | 187 | Gly (G) - Asp (D) | Increased virus binding to alpha2-6 glycans | [66] |
|  | 190 | Glu (E) - Asp (D) | Specificity mammalian transmissibility | [20] |
|  | 222 | Asp (D) - Gly (G) | Increased pathogenicity | [45] |
|  | 225 | Gly (G) - Asp (D) | Receptor-binding specificity | [11] |
|  | 226 | Gln (Q) - Leu (L) | Receptor-binding specificity | [11] |
|  | 228 | Gly (G) - Ser (S) | Associated with virulence | [49] |
| NA<sup>1</sup> | 223 | Thr (T) - Ile (I) | Associated with virulence | [6] |
|  | 275 | His (H) - Tyr (Y) |  | [27] |
| PB2 | 357 | His (H) - Asn (N) | Increased polymerase activity | [10] |
|  | 591 | Gln (Q) - Lys (K) | Enhance viral replication | [10] |
|  | 627 | Glu (E) - Lys (K) | Polymerase activity | [60] |
|  | 701 | Asp (D) - Asn (N) | Mammalian transmissibility | [22] |
| PA | 35 | Phe (F) - Leu (L) | Associated with pandemics | [45] |
|  | 97 | Thr (T) - Ile (I) | Associated with virulence | [51] |
|  | 224 | Ser (S) - Pro (P) | Increased polymerase activity | [50] |
|  | 383 | Asn (N) - Asp (D) |  |  |
| NS1 | 92 | Asp (D) - Glu (E) | Associated with virulence | [44] |

<sup>1</sup> Positions are based on H1 and N1 numbering system [4].

The mutations at different sites or segments may have distinct contributions for the virulence prediction. For example, it is known that the mutations at HA protein are more responsible for the viral activities, changing the receptor-binding specificity or increasing pathogenicity. To fully make use of these prior knowledge, we first obtain the prior mutation information 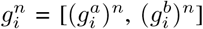 of sequence sample *n* at position *i*, and the corresponding label is 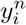. The feature on amino acids in certain sites can be defined as follows:

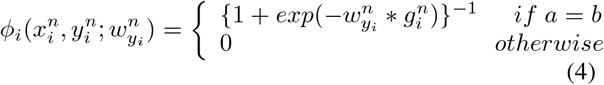

where 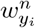 is the leaned parameter to represent the various effect of mutations at different proteins and sites, 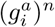 represents the input residue information at site *i* of sample *n* and 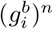 is the prior known mutated residue at the same site. To simplify the process, we only consider the mutations at different positions regardless of the proteins they occur. The value of *ϕ*_*i*_ determines the constraint feature at a certain site, illustrating the mutation information of input data. For most of the sequences, there is usually more than one mutation in the genomic sequence. The more mutations occurred, the more virulent it could be since the viruses may escape from the immune system and cause severe symptoms in humans. Therefore, we implement similar procedures to other prior mutation sites based on Table 1 to extract all the constraint features from viral mutations.

#### Reassortment probability

Reassortment is another key mechanism in the viral evolution, resulting in the emergence of novel strains. Different combinations of influenza segments from distinct parent strains could produce plenty of descendant strains. The reassortant strains have been responsible for several pandemics since the 19th century and have infected numerous people with severe disease and even death. It was suggested that the virulence of new viruses might increase after reassortment. Therefore, we assume if the influenza strains have been detected to be reassortant, then the probability of being virulent is higher than those that are non-reassortant. To obtain the reassortment information of input genomes, we employed a computational method named HopPER [67] to obtain the influenza reassortment probabilities through host tropism prediction. This model applied a kernel perspective on host probability estimation by random forest on individual sequence in the genome and integrated the results to calculate genome reassortment probability. We directly use *REprob*(*x*) to represent the function to compute reassortment probability of genome *x* based on HopPER. Thus, the constraint feature of reassortment information is formulated in Eq. (5), where *x*^*n*^ is the *n*-th input genome sample and *y*^*n*^ is the corresponding virulence label.

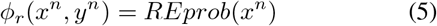

### 2.5 Model construction

To tackle the issue of accurately predicting influenza virulence with interpretation, we employed the posterior regularization technique [12]. The flowchart in Figure 2 describes the process of incorporating prior viral knowledge into the basic predictive model.

**Figure 2:**
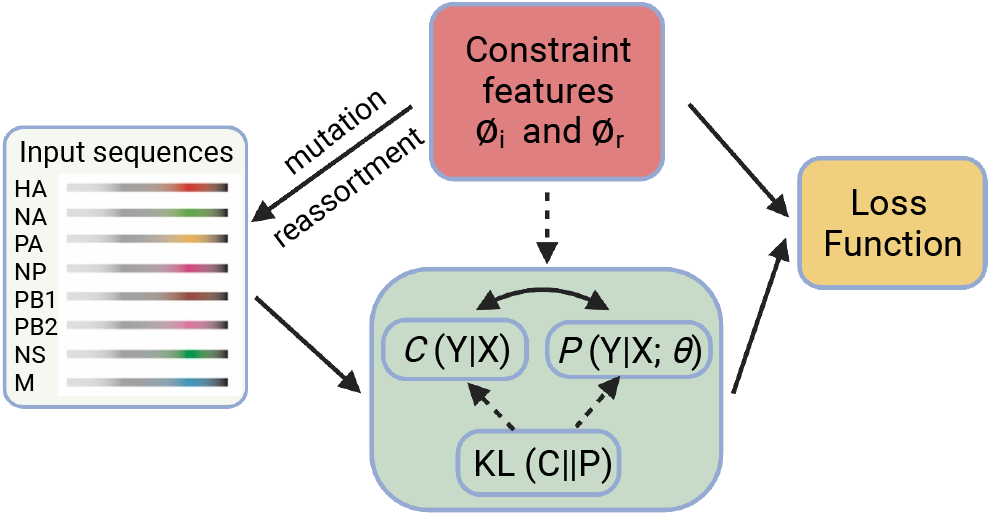
The flowchart of incorporating prior knowledge into basic predictive model.

Given the input data *X*^*n*^=[*x*^1^, *x*^2^, …, *x*^*n*^] that each sample is the concatenated genome containing all 8 segments, and the label vector is denoted as *Y* ^*n*^=[*y*^1^, *y*^2^, …, *y*^*n*^]. The prediction model can be adopted to acquire the predictive probability vector *Ŷ* ^*n*^ = P(*Y* ^*n*^|*X*^*n*^; *θ*). The contribution of the proposed framework is to combine prior viral information into the existing model by utilizing the desired distribution with posterior regularization. The main goal of the posterior regularization is to confine the space of the model posteriors using prior information to guide the model towards desired parameter distributions [12] that has been widely applied to the prediction tasks [18, 32]. In this case, posterior information is denoted with sets *C* of allowed distributions over hidden variables *Y*. Adapting from the risk prediction on electronic health records [32], we define *c*(*y*^*n*^) as the desired distribution of sample *n*, and the posterior regularized loss function can be formulated as

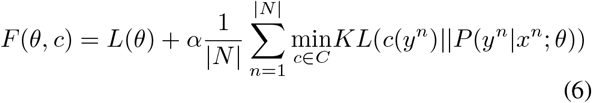

where *L*(*θ*) is the raw objective function of virulence prediction indicating average cross-entropy, and *α* is a hyperparamter that makes a balance between the posterior regularization and the loss of the predictive model. Kullback-Leibler divergence [38] (KL(·||·)) measures the difference between *c*(*y*^*n*^) and the posterior distribution *P* (*y*^*n*^|*x*^*n*^; *θ*) of the predictive model. *N* is the set of genomic sequences and |*N* | denotes the number of input sequences, where *n* ∈ |*N*|. The set of constraints for posterior information is represented as *C* defined by:

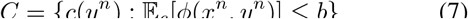

where *ϕ*(*x*^*n*^, *y*^*n*^) is the constraint features on sample *n* and *b* denotes the bounds on the desired expected values of constraint features. To obtain the optimized parameters for virulence prediction model, we use *J* (*θ, τ, w*) to represent the objective function that can be reformulated as:

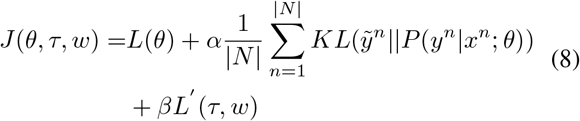

where the desired distribution 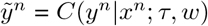, which encodes prior mutation and reassortment information. The mathematical equation of 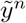 is defined below:

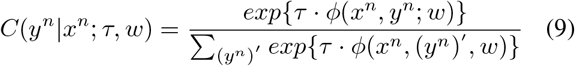

where *τ* is the updated confidence matrix for different categories of constraint features. The parameter *w* is added to successfully distinguish the difference among multiple pieces of the same categorical prior information. *β* is the hyperparameter and *L′* (*τ, w*) stands for the average cross entropy between the ground truth *y*^*n*^ and desired distribution 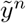. The equation of *L′* (*τ, w*) can be found:

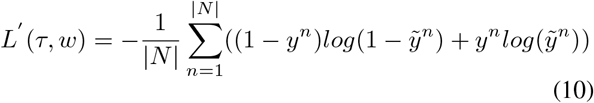

From these equations, we can observe that the proposed method is a general framework that integrates prior knowledge into the predictive model. The prior knowledge in this work includes viral mutations and reassortments, but more information can be merged into the framework. Furthermore, the goal of the proposed model is to minimize the objective function by learning a set of parameters and hyperparameters. Combining the effect of prior viral knowledge and updated parameters, we use the following equation to predict the virulence of influenza virus:

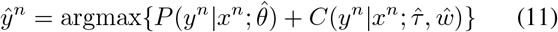

## 3 Experimental setup

### 3.1 Baseline approaches

To evaluate the performance of the proposed framework for virulence prediction, we implemented several types of baselines. The first type is traditional classification approaches that we compare ViPal with logistic regression (LR), naïve bayes (NB), K-nearest neighbor (KNN) and support vector machine (SVM). The detailed information of these classifiers are shown in Supplementary Materials S3. The second type is using deep learning-based approaches. Four variants of CNN architecture AlexNet [25], VGG-16 (Visual Geometry Group) [46], ResNet-50 [16] and SqueezeNet [19]. The proposed framework is based on the deep learning models by incorporating prior knowledge with the posterior regularization technique. They are denoted as AlexNet*, VGG-16*, ResNet-50* and SqueezeNet*. The settings of the proposed models are the same as deep learning-based baselines.

### 3.2 Implementation and evaluation

All the approaches are implemented with Scikit-learn [37] and PyTorch [35]. The processed data is randomly divided into training and testing samples in a ratio of 0.8:0.2. In the training process, 5-fold cross validation is performed and the testing set is used to select the values of the parameters. For traditional machine learning methods, the parameters are set to their default values. For all deep learning-based models, stochastic gradient descent is applied with a minimum batch size of 4 for optimization. The learning rate is 0.0001 and with 147 filter maps. The L1 regularization and drop-out (rate = 0.5) strategy are carried out for all deep learning approaches with 100 training epochs. Besides, we set *α* = 1 and *β* = 1 for all the proposed models. Lastly, we performed an interpretability analysis using SHAP (SHapley Additive exPlanation) values [31], a game-theoretic approach to model explanations, which can assist us in uncovering the most important constraint features for the prediction. The evaluation metrics consist of accuracy, precision, recall and F-score in all methods for virulence prediction.

## 4 Results and discussion

### 4.1 Performance comparison with baselines

We first investigated the two new hyperparameters *α* and *β* in Eq. (8). We defined *α, β* ∈ [0.1, 0.2, 0.3, 0.4, 0.5, 0.6, 0.7, 0.8, 0.9, 1.0, 2.0, 3.0]. The sensitivity analysis was implemented by changing one hyperparameter while fixing the other to track the variation of predictive performance. The results did not suggest a clear relation between the hyperparameters and the performance (Supplementary Materials S4). Therefore, we empirically set *α* = 1 and *β* = 1 for the subsequent experiments. Table 2 shows the results of all the methods on the processed influenza dataset for virulence prediction. We could observe that the proposed models displayed better performance than all other baselines in terms of most of the measures on the testing dataset. In more detail, the overall performance of KNN, LR, NB and SVM was slightly worse than the deep learning models, which suggested that leveraging deep learning techniques is more effective to deal with the high dimensional feature space issues. Among all the deep learning-based baselines, AlexNet achieved the best performance than SqueezeNet, VGG-16 and ResNet-50. However, we observed that the overall performance of deep learning baselines was slightly better than the proposed model. It is partially because DNN is constructing deep and complicated architecture. Since the size of the influenza dataset is relatively small with transformed features in high dimension, it is easily cause overfitting problems with deep neural network models. Vipal reduced this issue by adding prior viral information and the results showed that all the proposed approaches outperformed their corresponding baseline methods in the testing data. VGG-16* exhibited superior performance than AlexNet*, SqueezeNet* and ResNet-50*. These observations strongly confirmed that incorporating prior viral knowledge would help the prediction models to improve the performance. Moreover, it further demonstrated the efficiency of transforming prior knowledge into the prediction models by posterior regularization techniques.

**Table 2:** Comparative performance of virulence prediction on influenza dataset. Acc: Accuracy, Pre: Precision, Rec: Recall, F1: F-score.

| Model |  | Training |  |  |  | Testing |  |  |  |
| --- | --- | --- | --- | --- | --- | --- | --- | --- | --- |
|  |  | Acc | Pre | Rec | F1 | Acc | Pre | Rec | F1 |
| Traditional Classifier | KNN | 0.771 | 0.773 | 0.914 | 0.838 | 0.653 | 0.696 | 0.821 | 0.754 |
|  | SVM | 0.672 | 0.731 | 0.777 | 0.753 | 0.639 | 0.710 | 0.747 | 0.728 |
|  | NB | 0.533 | 0.658 | 0.540 | 0.581 | 0.531 | 0.710 | 0.611 | 0.657 |
|  | LR | 0.771 | 0.773 | 0.914 | 0.838 | 0.653 | 0.696 | 0.821 | 0.754 |
| Deep learning model | AlexNet | 0.769 | 0.767 | **0.916** | 0.835 | 0.694 | 0.743 | 0.833 | 0.786 |
|  | ResNet-50 | 0.799 | **0.924** | 0.746 | 0.825 | 0.602 | 0.671 | 0.803 | 0.731 |
|  | VGG-16 | 0.800 | 0.803 | 0.911 | **0.853** | 0.643 | 0.718 | 0.773 | 0.745 |
|  | SqueezeNet | 0.787 | 0.791 | 0.906 | 0.845 | 0.673 | 0.743 | 0.788 | 0.765 |
| Proposed Approach | AlexNet* | 0.748 | 0.813 | 0.796 | 0.804 | 0.776 | **0.816** | 0.886 | 0.849 |
|  | ResNet-50* | **0.826** | 0.857 | 0.892 | 0.874 | 0.755 | 0.793 | 0.793 | 0.793 |
|  | VGG-16* | 0.772 | 0.721 | 0.836 | 0.828 | **0.816** | **0.816** | **0.939** | **0.873** |
|  | SqueezeNet* | 0.760 | 0.811 | 0.836 | 0.823 | 0.776 | 0.765 | 0.897 | 0.825 |

### 4.2 Constraint feature analysis

To ensure the reliability and stability of such a framework, human needs to understand how these decisions are made. Figure 3 illustrated the SHAP values for top 10 constraint features by VGG-16* model. The constraint features in the y-axis are organized by their SHAP values (x-axis), which represented the importance of driving the prediction of the classifiers for virulence prediction. The variables were ranked in descending order and the SHAP values plot can further show the positive and negative associations of the predictors with the target. According to the results, we discovered that the mutations on site 627 of PB2 protein and site 92 of NS1 protein play more important roles than other constraint features. Besides, prior reassortment information also ranked on top among all the constraint features. Interestingly, some of the features were revealed to be negatively associated with the prediction model, e.g., site 187 on the HA protein. One explanation was that the insufficiency of training data hindered a comprehensive discovery of the relations between mutation and virulence. Nevertheless, this corroboration of the features learned by our method and highlighted by the SHAP analysis with the results from prior knowledge exploring influenza virulence gave additional credibility to previous findings.

**Figure 3:**
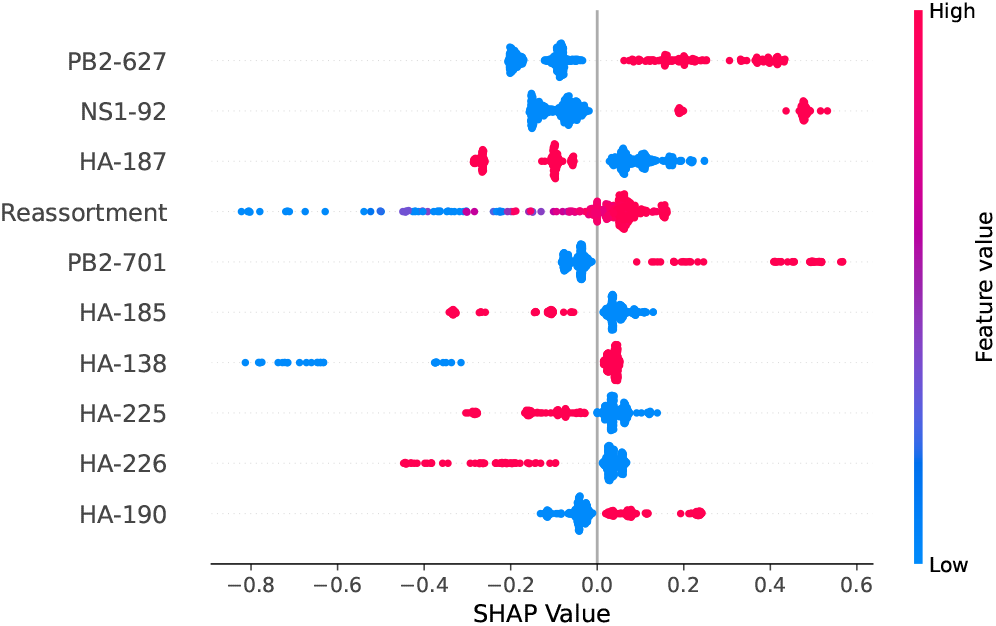
The plots of SHAP analysis showing the 10 most important constraint features by VGG-16* model for virulence prediction of influenza A virus.

### 4.3 Comparison with existing methods

We further compared our proposed framework with two existing methods for computational virulence prediction. Ivan et al. [21] applied three different rule-based approaches with additionally random forest for modeling. The model PART achieved the best performance. The other approach VirPreNet [62] developed a weighted ensemble convolutional neural network for virulence prediction of influenza A viruses based on all eight segments. Figure 4 presented predictive performance and computation cost between ViPal and the existing models. The prediction performance included accuracy, precision, recall. According to the results, VirPal obtained better outcomes in accuracy and precision and was comparable in recall compared with VirPreNet. Both methods remarkably outperformed PART. However, VirPreNet spent much more time training the model for limited enhancement as it constructed a weighted ensemble deep neural network. The input data of each genomic segment was fed to build an ensemble model as a base model. We cannot obtain the final prediction until all the eight ensemble models are built, followed by a linear layer. Our proposed model significantly reduced the computation time by taking advantage of the concatenation of different influenza proteins based on CTD descriptors. Interestingly, PART required the least computation time, but the results were not as competitive as the other two models. Thus, ViPal is more superior regarding the performance of prediction and computation cost and has the potential for virulence prediction when being trained on large datasets.

**Figure 4:**
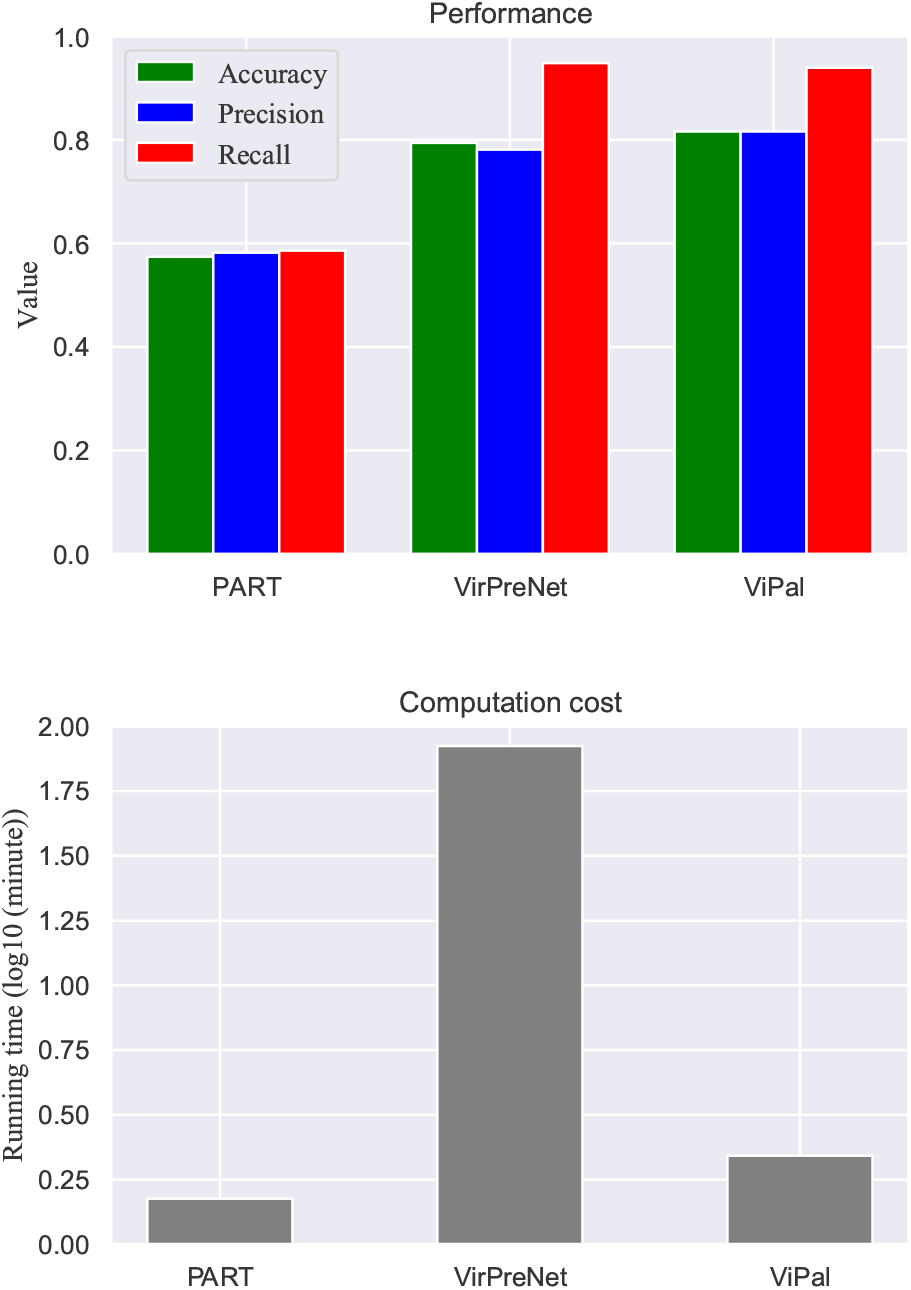
Comparison between ViPal and two existing models regarding their predictive performance and computation cost. X-axis is three different methods. Y-axis is the performance values at the top and running time (*log*_10_(*minute*)) in the bottom.

### 4.4 Model evaluation on individual subtypes

To validate the generalizability of the proposed framework, we conducted experiments on individual influenza subtypes for virulence prediction. The influenza dataset was divided into different categories based on their subtypes. As a result, we obtained 150 samples for H1N1, 42 samples for H3N2 and 153 samples for H5N1. The remaining samples were combined into one group labeled as “Others” due to the very few cases of other subtypes. Table 3 presented the predictive performance on the testing dataset of each category. Comparing our proposed models with their corresponding baselines without prior knowledge, we can observe an apparent increase in terms of most measures. The results demonstrated that even on the smaller dataset, incorporating prior viral knowledge can still strengthen the predictive performance. Among all the methods, VGG-16* suggested a superior performance over other models in H1N1, H5N1 and other subtypes. It was also found that, overall, VGG-16* made greater improvement over VGG-16 in comparison to AlexNet*/AlexNet, ResNet-50*/ResNet-50 and SqueezeNet*/SqueezeNet. Surprisingly, SqueezeNet* performed perfectly on virulence prediction on subtype H3N2 that all the cases were correctly predicted. This is probably because the distribution of the H3N2 dataset is well-matched with the model. In summary, according to Table 2 and 3, we can safely ascertain the effectiveness of incorporating prior viral knowledge into existing prediction models for the improvement of predictive performance.

**Table 3:** Performance of virulence prediction on individual subtypes of influenza A viruses. Acc: Accuracy, Pre: Precision, Rec: Recall, F1: F-score.

| Subtype | Model | Acc | Pre | Rec | F1 |
| --- | --- | --- | --- | --- | --- |
| H1N1 | AlexNet | 0.733 | 0.775 | 0.912 | 0.838 |
|  | ResNet-50 | 0.644 | 0.725 | 0.853 | 0.784 |
|  | VGG-16 | 0.667 | 0.741 | 0.870 | 0.800 |
|  | SqueezeNet | 0.689 | 0.738 | 0.912 | 0.816 |
|  | AlexNet* | 0.756 | 0.738 | **1.000** | 0.849 |
|  | ResNet-50* | 0.689 | 0.758 | 0.806 | 0.781 |
|  | VGG-16* | **0.767** | **0.786** | 0.957 | **0.863** |
|  | SqueezeNet* | 0.733 | 0.721 | 1.000 | 0.849 |
| H3N2 | AlexNet | 0.769 | 1.000 | 0.700 | 0.824 |
|  | ResNet-50 | 0.692 | 0.875 | 0.700 | 0.778 |
|  | VGG-16 | 0.667 | 0.714 | 0.833 | 0.769 |
|  | SqueezeNet | 0.889 | 1.000 | 0.833 | 0.909 |
|  | AlexNet* | 0.923 | 1.000 | 0.857 | 0.923 |
|  | ResNet-50* | 0.769 | 1.000 | 0.571 | 0.727 |
|  | VGG-16* | 0.889 | 1.000 | 0.833 | 0.909 |
|  | SqueezeNet* | **1.000** | **1.000** | **1.000** | **1.000** |
| H5N1 | AlexNet | 0.677 | 0.700 | 0.955 | 0.808 |
|  | ResNet-50 | 0.565 | 0.679 | 0.633 | 0.655 |
|  | VGG-16 | 0.710 | 0.710 | **1.000** | 0.83 |
|  | SqueezeNet | 0.645 | 0.762 | 0.727 | 0.744 |
|  | AlexNet* | 0.710 | 0.810 | 0.773 | **0.891** |
|  | ResNet-50* | 0.630 | 0.667 | 0.786 | 0.721 |
|  | VGG-16* | **0.742** | **0.818** | 0.818 | 0.818 |
|  | SqueezeNet* | 0.677 | 0.800 | 0.727 | 0.762 |
| Others | AlexNet | 0.605 | 0.621 | 0.750 | 0.679 |
|  | ResNet-50 | 0.558 | 0.586 | 0.708 | 0.642 |
|  | VGG-16 | 0.655 | 0.684 | 0.765 | 0.722 |
|  | SqueezeNet | 0.698 | 0.704 | 0.792 | 0.745 |
|  | AlexNet* | 0.837 | 0.920 | 0.821 | 0.868 |
|  | ResNet-50* | 0.814 | 0.917 | 0.786 | 0.846 |
|  | VGG-16* | **0.897** | **0.938** | **0.882** | **0.909** |
|  | SqueezeNet* | 0.791 | 0.828 | 0.857 | 0.842 |

## 5 Conclusion

In this paper, we proposed a general framework ViPal for the virulence prediction of influenza A viruses through machine learning methods using all influenza segments, integrating prior viral knowledge into all the existing predictive models to improve the predictive performance. Specifically, we employed four traditional machine learning models and four state-of-the-art deep learning architectures as the basic predictive models. To model the prior influenza information into the model, we leveraged the posterior regularization techniques. The discrete and heterogeneous prior knowledge was converted into continuous values by modeling the posterior distribution as the constraint feature set. Experimental results indicate that ViPal exceeds the baselines and two off-the-shelf approaches. Furthermore, the interpretability analysis revealed the importance of different prior constraint features on the model, revealing virulence variation due to the mutation and reassortment.

The future work can be oriented in several directions. Firstly, we use mutation and reassortment as prior knowledge to build the model, while some other factors will also impact the influenza virulence, e.g., host-pathogen interaction and environmental influence. Taking more and diverse prior information into consideration could further improve the predictive performance and potentially better understand influenza virulence when constructing the model. Secondly, we only focus on influenza virulence prediction in this work, whereas it can be extended to other viruses such as the Ebola virus and coronavirus. Further experiments are required to validate. Thirdly, to demonstrate the flexibility of this method when applied to large datasets, we will search for more qualified influenza samples through literature review or other public databases for external testing. We believe that ViPal has the ability to handle a larger dataset and the algorithms proposed here can even have the potential to predict the progression or lethality in other diseases with appropriate adjustment.

