## Supplementary Materials S for "ViPal: A Framework for Virulence Prediction of Influenza Viruses with Prior Viral Knowledge Using Genomic Sequences"

S1. The influenza datasets used in this study can be found at the following link:

https://github.com/Rayin-saber/ViPal/tree/main/data

S2. Table: The division of amino acid groups based on physicochemical properties and amino acid indices.

| Attributes | Group 1 | Group 2 | Group 3 |
|---|---|---|---|
| Hydrophobicity | Polar<br>Q, E, R, K, D, N | Neutral<br>G,P, H, A, S, T, Y | Hydrophobic<br>C, V, F, L, I, M, W |
| Polarizability | 0-1.08<br>S, D, G, A, T | 0.128-0.186<br>C, Q, I, P, N, V, E, L | 0.219- 0.409<br>Y, M, K, R, H, F, W |
| Normalized Van der Waals | 0-2.78<br>S, C, G, A, T, P, D | 2.95-4.0<br>E, Q, N, V, I, L | 4.0-8.1<br>K, F, M, H, R, Y, W |
| Polarity | 4.9-6.2<br>W, C, L, I, F, M, V, Y | 8.0-9.2<br>T, G, P, A, S | 10.4-13.0<br>K, N, H, Q, R, E, D |
| Solvent Accessibility | Buried<br>A, I, F, C, G, L, V, W | Exposed<br>R, K, Q, E, N, D | Intermediate<br>M, S, P, T, H, Y |
| Secondary Structure | Helix<br>E, A, L, M, Q, K, R, H | Strand<br>V, I, Y, C, W, F, T | Coil<br>G, N, P, S, D |
| Charge | Positive<br>K, R | Neutral<br>A, N, C, Q, G, H, I, L, M, F, P, S, T, W, Y, V | Negative<br>D, E |

S3. The parameter setting for traditional machine learning classifiers.

Logistic Regression: penalty='L2', tol =0.0001, c=1.0, intercept_scaling=1, class_weight=None, max_iter=100

K-nearest neighbor: n_neighbors=5, weights='uniform', algorithm='auto', leaf_size=30, p=2, metric='minkowski', metric_params=None, n_jobs=None

Support vector machinne: C=1.0, kernel='rbf', degree=3, gamma='scale', coef0=0.0, shrinking=True, probability=False, tol=0.001, cache_size=200, class_weight=None, verbose=False, max_iter=-1, decision_function_shape='ovr', break_ties=False,

Naïve bayes: alpha=1.0, binarize=0.0, fit_prior=True, class_prior=None

S4. Table: Performance on different values of hyperparameters α and β on testing data for virulence prediction with ResNet-50*.

| α (β=1) | Testing data | | | | |
|---|---|---|---|---|---|
| | Accuracy | Precision | Recall | F-score | AUC |
| **0** | 0.745 | 0.824 | 0.836 | 0.830 | 0.512 |
| **0.1** | 0.745 | 0.824 | 0.836 | 0.830 | 0.515 |
| **0.2** | 0.745 | 0.824 | 0.836 | 0.830 | 0.524 |
| **0.3** | 0.745 | 0.824 | 0.836 | 0.830 | 0.528 |
| **0.4** | 0.745 | 0.824 | 0.836 | 0.830 | 0.528 |
| **0.5** | 0.745 | 0.824 | 0.836 | 0.830 | 0.527 |
| **0.6** | 0.745 | 0.824 | 0.836 | 0.830 | 0.529 |
| **0.7** | 0.745 | 0.824 | 0.836 | 0.830 | 0.529 |
| **0.8** | 0.735 | 0.822 | 0.822 | 0.822 | 0.530 |
| **0.9** | 0.745 | 0.824 | 0.836 | 0.830 | 0.528 |
| **1** | 0.735 | 0.822 | 0.822 | 0.822 | 0.545 |
| **2** | 0.735 | 0.822 | 0.822 | 0.822 | 0.544 |
| **3** | 0.735 | 0.822 | 0.822 | 0.822 | 0.550 |

| β (α=1) | Testing data | | | | |
|---|---|---|---|---|---|
| | Accuracy | Precision | Recall | F-score | AUC |
| **0** | 0.755 | 0.818 | 0.863 | 0.840 | 0.608 |
| **0.1** | 0.765 | 0.838 | 0.849 | 0.844 | 0.464 |
| **0.2** | 0.786 | 0.842 | 0.877 | 0.859 | 0.561 |
| **0.3** | 0.745 | 0.808 | 0.863 | 0.834 | 0.531 |
| **0.4** | 0.653 | 0.831 | 0.671 | 0.742 | 0.506 |
| **0.5** | 0.755 | 0.836 | 0.836 | 0.836 | 0.610 |
| **0.6** | 0.724 | 0.838 | 0.781 | 0.809 | 0.582 |
| **0.7** | 0.745 | 0.824 | 0.836 | 0.830 | 0.615 |
| **0.8** | 0.704 | 0.789 | 0.822 | 0.805 | 0.597 |
| **0.9** | 0.765 | 0.821 | 0.877 | 0.848 | 0.649 |
| **1** | 0.745 | 0.824 | 0.836 | 0.830 | 0.693 |
| **2** | 0.714 | 0.800 | 0.822 | 0.811 | 0.580 |
| **3** | 0.724 | 0.838 | 0.781 | 0.809 | 0.664 |
